# *Fragaria ananassa DAM4* expression correlates with vegetative growth during semi-dormancy breaking

**DOI:** 10.1101/2025.07.10.664089

**Authors:** Stephan David, Philippe Kersten, Xiren Cao, Leo F.M. Marcelis, Julian C. Verdonk

## Abstract

*Dormancy-Associated MADS-BOX (DAM)3* and *DAM4* have been described as potential regulators of winter dormancy in cultivated strawberry (*Fragaria* x *ananassa* Duch.). These genes are upregulated under short day conditions and downregulated under chilling conditions. The aim of the current work is to test the hypothesis that the expression of *DAM3* and *DAM4* correlates negatively with vegetative growth during dormancy induction and breaking. The expression of *DAM3* and *DAM4* as well as plant morphology and physiology were analyzed during a period of semi-dormancy induction and breaking. Furthermore, *DAM3* and *DAM4* expression was compared between cultivars with different chill requirements. Lastly, the *DAM1, DAM2, DAM3*, and *DAM4* expression of concurrently growing summer and winter leaves was compared at multiple stages of dormancy-breaking. *DAM3* and *DAM4* expression negatively correlated with leaf area and petiole length. We conclude that *DAM3* and *DAM4* expression patterns are consistent with that of regulators of semi-dormancy. *DAM4* expression in particular correlates strongly with vegetative growth during semi-dormancy breaking. *DAM3* or *DAM4* expression were not found to correlate with the cultivar-specific chill requirements. Lastly, no relevant differences in *DAM* expression were found between summer and winter leaves.

## INTRODUCTION

Many climates have a period of environmental conditions that are unfavorable to plant growth, such as a cold or a dry season. Annual plants can bridge this period as a seed and sprout once conditions are more favorable for growth. Perennials require a developmental strategy in order to survive the adverse conditions until the next growing season. Typically, this strategy consists of a period of reduced or halted growth, referred to as dormancy, to protect meristematic tissues from harsh conditions.

In cultivated strawberry (*Fragaria* x *ananassa* Duch.) and its wild ancestor woodland strawberry (*F. vesca*), induction of winter dormancy does not lead to cessation of meristem growth. This has prompted the term semi-dormancy^1,2^. Between the summer and winter, leaf development changes, resulting in a different phenotype. Winter leaves are smaller, thicker, have shorter petioles, and contain more chlorophyll per unit area than their summer counterparts, giving semi-dormant plants a stunted appearance^3^. The difference in phenotype between the summer and winter is known as seasonal dimorphism. These adaptations allow winter leaves to survive the winter cold and drought, whereas summer leaves senesce in sufficiently cold temperatures. The development of winter-hardy leaves allows for photosynthesis in the autumn and mild winter conditions, extending the growing season.

The induction of semi-dormancy is a quantitative process, which, simultaneously with flower initiation, occurs under short day conditions^2^. Similar to many species which exhibit a form of winter-dormancy, exposure to cold breaks semi-dormancy and reverts vegetative growth to the summer phenotype^4^. Interestingly, exposure to sufficient amounts of chilling results in inhibition of subsequent flower initiation, which prevents induction of generative meristem under exposure to short photoperiod in early spring^5^.

The transition from annuals to perennials in Rosaceous plants is speculated to have been facilitated by the diversification of SHORT VEGETATIVE PHASE (SVP), a protein family of MADS-box transcription factors^6^. DORMANCY ASSOCIATED MADS-BOX (DAM), a subfamily of the SVP family, was first described after a knock-out of several *DAM* genes was found to be responsible for the naturally occurring peach (*Prunus persica*) *evergrowing* mutant^7,8^. This mutant does not exhibit winter dormancy and retains activity of terminal buds when exposed to conditions which are normally dormancy-inductive. Several genes have been identified in strawberry as homologs of the peach *DAM* genes^9,10^. Analysis of the protein structure and phylogeny showed that these are MIKC-type MADS-box proteins and that they exist in cultivated strawberry in pairs of two highly similar proteins^10^. Previously, we showed that *DAM3* and *DAM4* are up- and downregulated in strawberry plants when exposed to short day conditions and chilling respectively, alluding to their role as dormancy regulators^10^. We also found potential differential expression of *DAM1* and *DAM2* between summer leaves and winter leaves. Two cultivars with different chill requirements were included in this previous work, but no clear difference in *DAM* expression was found between them.

Based on our earlier findings, we hypothesize that *DAM3* and *DAM4* expression correlates negatively with vegetative growth during semi-dormancy induction and breaking. We hypothesize that there is no correlation between the chill requirement of different cultivars and *DAM3* and *DAM4* expression. Lastly, we hypothesize that with increased accumulation of chilling hours, the expression of *DAM1* and *DAM2* increases in summer leaves compared to winter leaves.

The semi-dormancy status of the plants was not considered in the analysis of the previous work^10^. In the current work, we aim to examine the correlation between vegetative growth as influenced by the semi-dormancy status of the plant, and *DAM3* and *DAM4* expression. We also explore the relationship between the chill requirement of a cultivar and the expression of *DAM3* and *DAM4*. Lastly, we wanted to find out whether there is differential expression of any of the *DAM* genes between concurrently growing winter and summer leaves. To this end, four strawberry cultivars were grown in a greenhouse under controlled conditions, designed to induce, inhibit or break semi-dormancy. Plant morphology and *DAM* expression were measured and analyzed.

## MATERIAL AND METHODS

### Plant material

Four seasonal flowering strawberry (*F. ananassa*) cultivars with different chill requirements from breeding company Fresh Forward (Huissen, The Netherlands) were used (Table 1). The plants were propagated from runners in plugs containing coconut fiber and peat substrate. The plug plants were grown under natural long day conditions in the greenhouse facilities of Fresh Forward (51.9°N, 5.9°E.) and frequently irrigated to promote rooting.

**Table 1.**
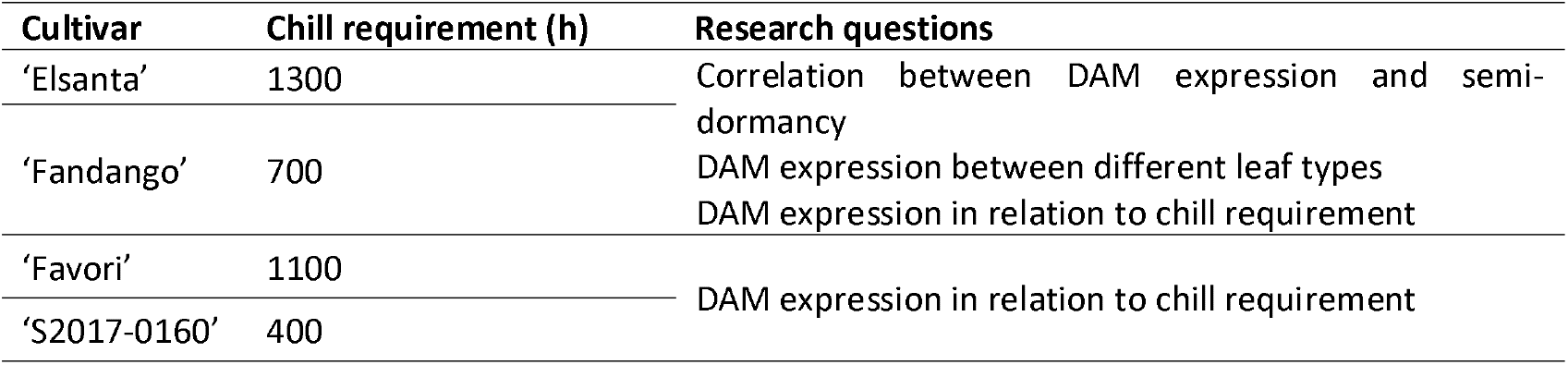
list of cultivars. Chill requirement denotes approximate number of hours needed at 7°C or less to break semi-dormancy for optimal vegetative growth, as indicated by the breeder. Research questions for each cultivar denote in which part of the experiment they were used.

### Plant cultivation

#### Starting phase

On August 29, 2023, well-rooted plug plants, containing 2-4 leaves, were transferred to the greenhouse and grown under natural light for six days. On September 4, two different growing conditions were created by activating supplementary lights in two compartments of the greenhouse: short day (12 h supplementary light) and far-red extended (12 h supplementary light extended to 16 h with far-red) (Figure 1). On the same day, 220 plants of the cultivars ‘Elsanta’ and ‘Fandango’ were placed in the short day compartment and 70 plants were placed in the far-red extended compartment. Of the cultivars ‘S2017-0160’ and ‘Favori’, 56 plants were placed in the short day compartment and none in the far-red extended.

**Figure 1.**
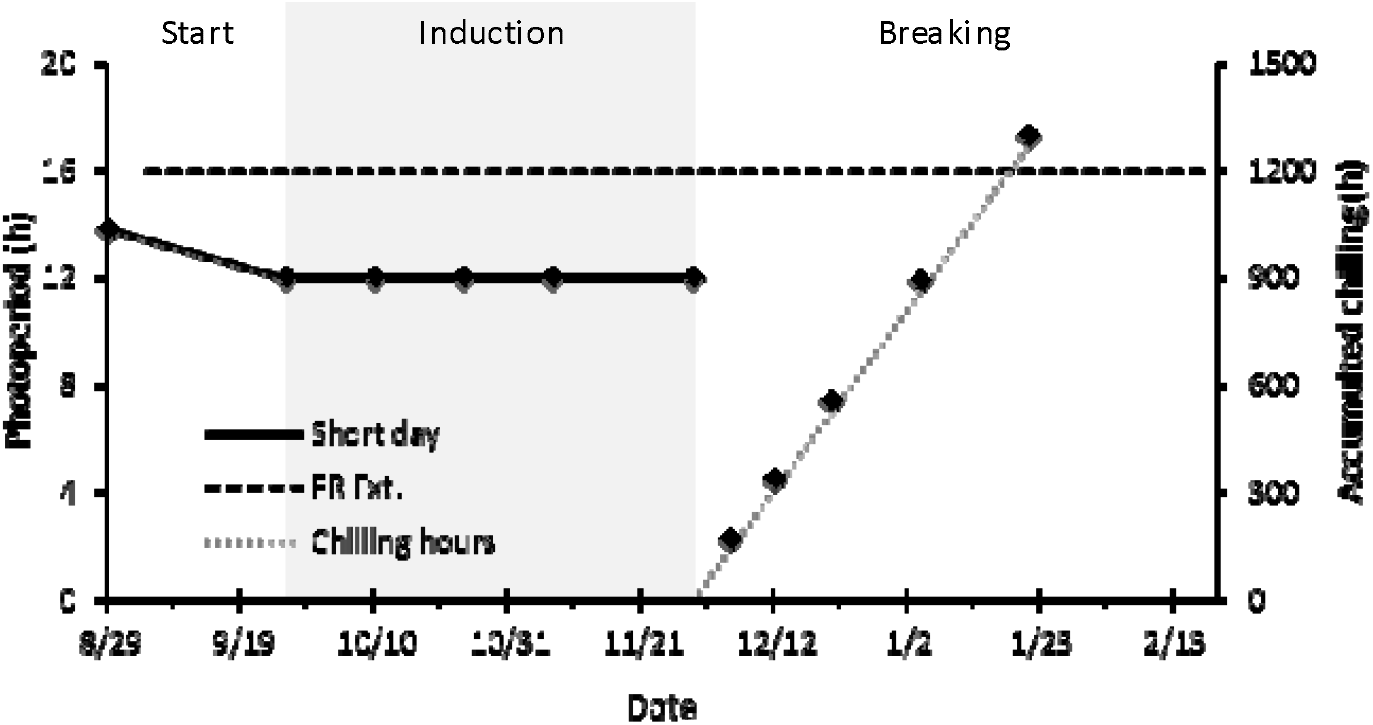
photoperiods in the short day and far-red extended (FR Ext.) compartments (primary y-axis) and accumulated chilling hours (secondary y-axis) during each experimental phase. Diamond markers on the far-red extended and chilling hours lines indicate dates on which leaf samples were taken and plants were forced. The grey background indicates the induction phase of the experiment.

#### Dormancy induction phase

On September 26, the natural photoperiod decreased to under 12 hours (11 h 58 m, Figure 1). Because of the supplemental lighting, the photoperiod in the short day compartment was now 12 h, and in the far-red extended compartment 16 h. We considered this the start of the dormancy induction phase, which lasted until November 29 (Figure 2). On November 29, the plants in the short day treatment had been exposed to a 12 hour photoperiod for 64 days.

**Figure 2.**
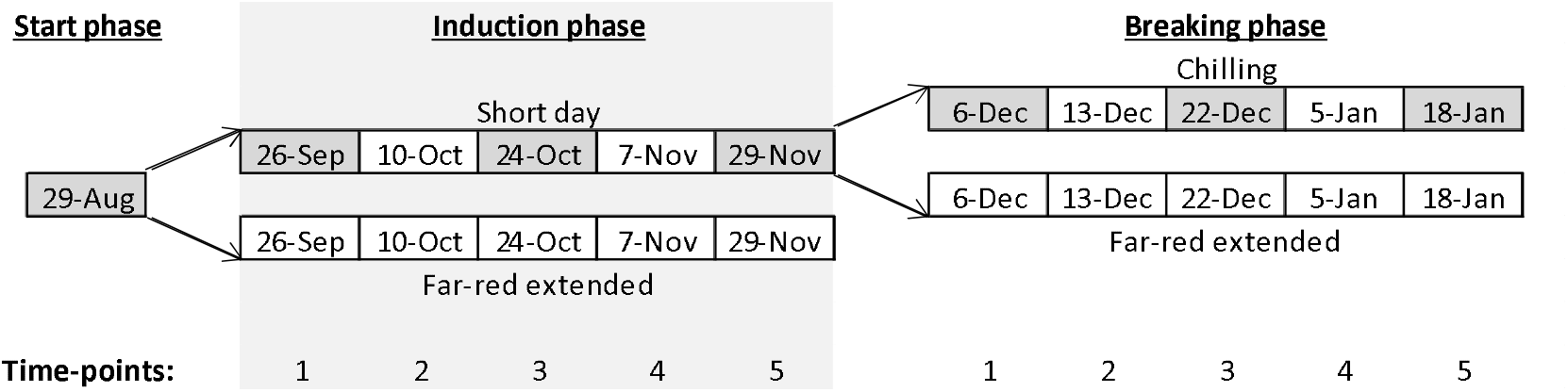
schematic overview of the experiment. Arrows indicate the transfer of plants from one phase to the next. Each block indicates a date on which leaf samples were taken and plants were potted and forced for four weeks. Cultivars ‘Elsanta’ and ‘Fandango’ were included in all treatments, time-points and measurements. Cultivars ‘Favori’ and ‘S2017-0160’ were only included in the short day and chilling treatments and leaf samples were only taken in the start phase, and on time points 1, 3, and 5 of the induction and breaking phase (marked in grey).

#### Dormancy breaking phase

On November 29, of the cultivars ‘Elsanta’ and ‘Fandango’, 80 plants from the short day compartment were transferred to the chilling treatment (2° C, complete darkness) and 70 were placed in the far-red extended treatment until the 18^th^ of January 2024 (Figure 2). All plants of the cultivars ‘S2017-0160’ and ‘Favori’ were transferred to the chilling treatment.

#### Forcing periods

On each time-point of each of the three phases of the experiment (Figure 2), eight plug plants of the cultivars ‘Elsanta’ and ‘Fandango’ from each treatment are transplanted in larger pots. These transplanted plants are grown in the far-red extended compartment for four weeks to stimulate vegetative growth. We refer to these four weeks as the forcing period.

### Growing conditions

In the short day treatment, plants received natural light supplemented with assimilation lamps (SON-T, Philips, the Netherlands, ±100 µmol·m^−2^·s^−1^ at plant level) from 6:00 until 18:00 (Supplement 1). In the far-red extended treatment, plants received the same hours of assimilation light, with the addition of low intensity light with a high far-red to red ratio (GreenPower LED Flowering Lamp Gen. 2.1, Philips, the Netherlands, ±10 µmol·m^−2^·s^−1^ at plant level) from 4:30 until 20:30 (Figure 1)(Supplement 2). This light was used in the far-red extended compartment to elicit a long day developmental response while minimally increasing the daily light integral. Because the natural photoperiod was longer than 12 h during the starting phase, the actual photoperiod in the short day compartment ranged from 13 h 48 min at the start of the starting phase to 12 h 2 m at the end. Short day and far-red extended treatments took place in the same greenhouse. Opaque plastic sheets reaching from the ceiling to the floor were hung around the far-red extended treatment to create two compartments and prevent light pollution. Temperature and humidity were measured with data loggers (Onset, HOBO MX2300) every 15 minutes from September 6 until the end of the experiment (Figure 3). The chilling treatment consisted of moving the plants from the greenhouse to cooling facilities at 2°C in complete darkness.

**Figure 3.**
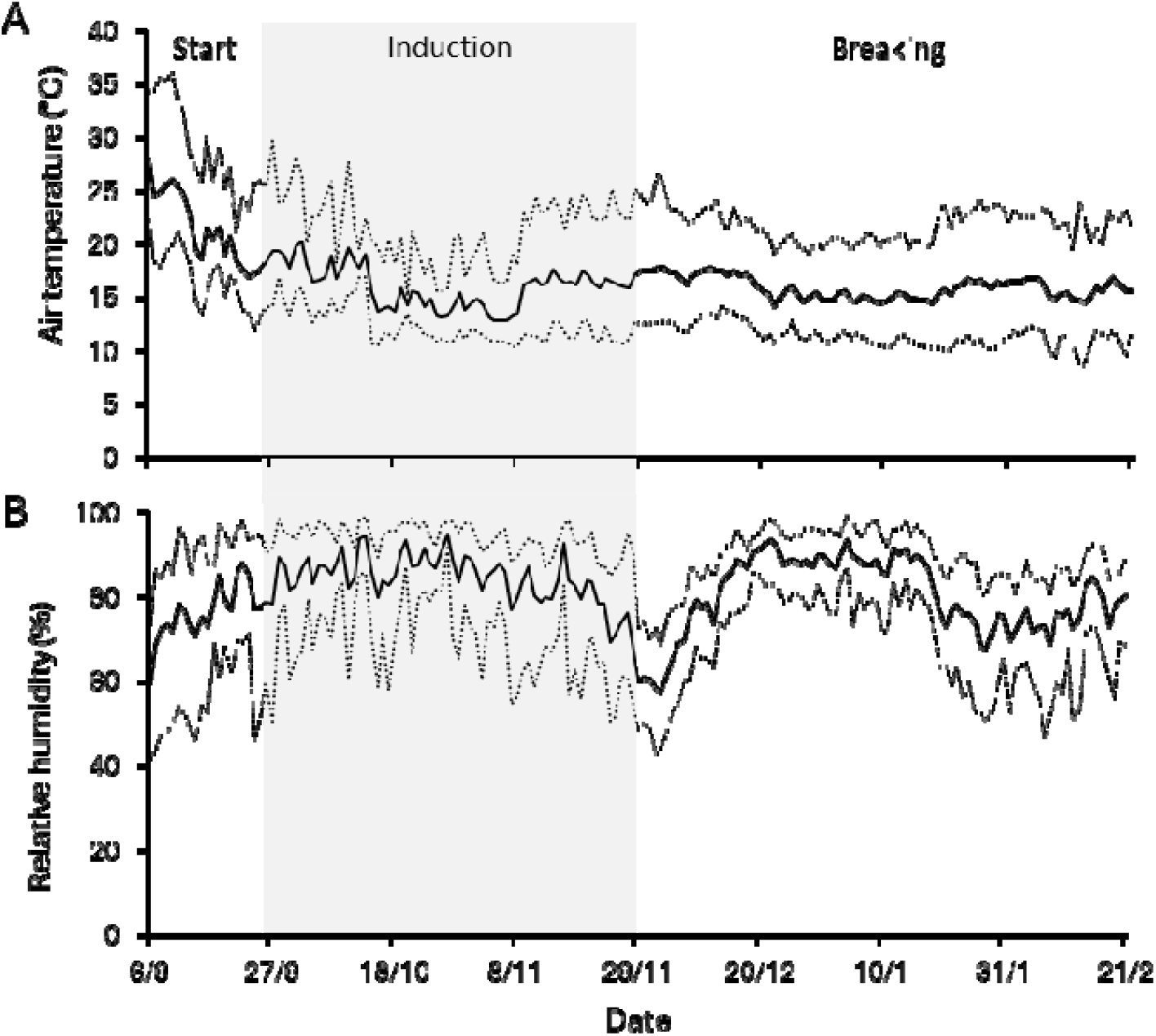
time course of daily (A) temperature and (B) relative humidity. Dotted lines denote the daily minimum and maximum values. Solid lines denote the daily average values. Air temperature and relative humidity were similar in both greenhouse compartments that were used in the experiment. The area marked in grey indicates the induction-phase.

### Correlation between *DAM* expression and semi-dormancy

On eleven time points, divided over the duration of the experiment, eight ‘Elsanta’ and ‘Fandango’ plug plants were taken from the short day or chilling, and far-red extended treatment for tissue sampling (Figure 1, Figure 2). Additionally, eight (short day and chill treatment) or six (far-red extended treatment) plug plants of the same cultivars were transferred to 5 liter pots with peat/coconut/perlite substrate (40:40:20 ratio), two plants per pot. All potted plants were placed in the far-red extended compartment to stimulate vegetative growth (forcing). Each leaf (with petiole of 1 cm or longer) was labeled and runners and flowers were removed. After four weeks of forcing, morphological and physiological measurements were taken. During the induction phase, after tissue sampling, the meristems were analyzed by dissection of the plants to determine the number of crowns per plant, number of meristem positions per plant, and meristem status (generative, vegetative or runner).

### *DAM* expression in relation to chill requirement

On seven time points, leaf tissue samples were taken from eight ‘Favori’ and ‘S2017-0160’ plants (Figure 2), after which these plants were removed from the experiment. The *DAM3* and *DAM4* expression levels and pattern of all four cultivars was compared in relation to the chilling requirement (Table 1).

### *DAM* expression between different leaf types

During the breaking phase, after forcing for four weeks, the ‘Elsanta’ and ‘Fandango’ plants contained leaves that had emerged during the induction phase (winter leaves), as well as leaves which emerged during forcing after chilling (referred to as summer leaves). Leaf samples were taken from both leaf types on every time point of the breaking phase. The expression of *DAM1, DAM2, DAM3, and DAM4* was compared between these two leaf types.

### Morphological and physiological measurements

All measurements were done on completely unfolded leaves that had emerged during forcing. Of each leaf we measured: the petiole length; the length of the middle leaf lobe and the width of the left leaf lobe; chlorophyll content (Force-A, Dualex, Orsay, France). Flowers and runners were counted as well. Leaf area was calculated from the leaf lobe measurements^11^.

### Gene expression analysis

Plants from the chilling treatment were acclimated at room temperature for two hours before leaf sample collection. Of each plant, two leaf samples of 1 cm in diameter were taken from two leaves. Leaf samples from two plants were pooled together resulting in four biological replicates from each cultivar and treatment. Samples were flash frozen in liquid nitrogen and stored at −80°C until further processing. From the ground leaf samples RNA was extracted, reverse transcribed into cDNA and used for gene expression analysis.

### RNA extraction

Leaf material was finely ground in a ball mill grinder (Retsch, MM400, Haan, Germany). RNA was extracted from the plant samples using the CTAB method (adapted from Schultz et al. 1994) with lithium chloride precipitation. All centrifuge steps were done in a microcentrifuge at maximum speed. In a 2 mL tube, 750 µL extraction buffer (1.4M NaCl, 20 mM EDTA pH8, 100 mM TRIS pH8, 2% CTAB, 2% PVP-40, 1% β-mercaptoethanol, milli-Q water (MQ) to 750 µL) was added to ±50 mg of frozen leaf powder. Samples were incubated 15 min at 65°C. 750 µL chloroform was added and samples were centrifuged at max speed for 5 min. The supernatant was transferred to a tube with 500 µL isopropanol, homogenized and centrifuged at max speed for 5 min. Supernatant was removed and pellet was washed with 70% ethanol. The pellet was dissolved in 205 µL MQ and RNA was precipitated by adding 67 µL 8M LiCl and incubating for 30 min at −20° C. The RNA was pelleted by centrifugation at max speed for 30 min at 4° C. Pellet was washed once more with 70% ethanol, air dried for 5 minutes and dissolved in 50 µL MQ. RNA quantity and quality were assessed with spectrophotometric analysis and gel-electrophoresis.

### cDNA synthesis

cDNA was synthesized from RNA (Applied Biosystems, High-Capacity cDNA Reverse Transcription Kit, Cat 4368813, Waltham, MA, USA). In a PCR tube, 200 ng RNA was combined with 2 µL 10x reaction buffer, 2 µL random primer mix, 0.8 µL 10 mM dNTP mix, 1 µL RTase enzyme, and MQ to 20 µL. Reverse-transcriptase PCR was done in a thermal cycler (SensoQuest Labcycler Gradient Fuses, Göttingen, Germany) with the following program: 10 min at 25° C, 120 min at 37° C, 5 min at 85° C. Finally, cDNA was diluted by adding 80 µL MQ. Quality of the cDNA was assessed by PCR amplification of *ACTIN1* (Table 2) and gel-electrophoresis.

**Table 2.**
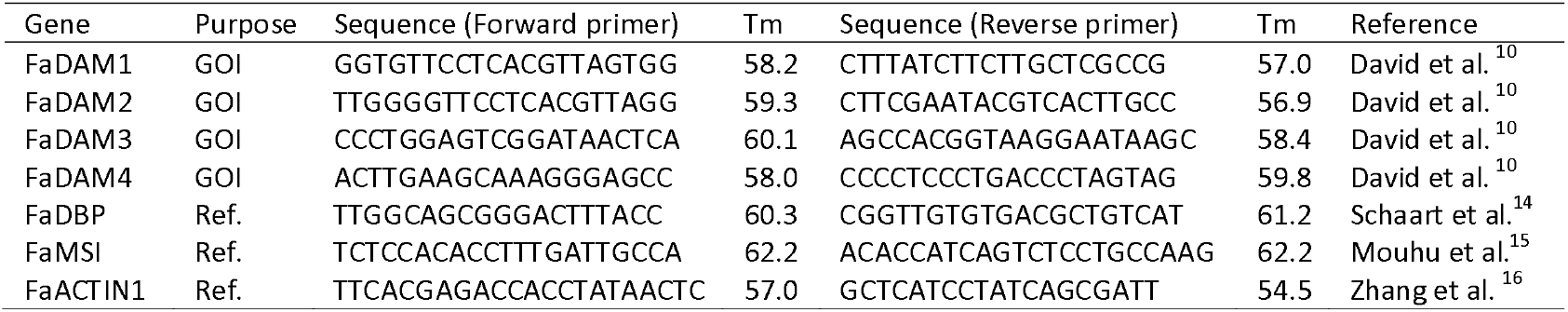
list of primers that were used for cDNA quality assessment with PCR and for expression analysis with qRT-PCR. GOI: gene of interest; Ref.: qRT-PCR reference gene.

### qRT-PCR

For each primer pair (Table 2), a primer mix was prepared (1 µL 10 µM forward primer, 1 µL µM reverse primer, 10 µL iQ SYBR Green Supermix (Bio-Rad, Hercules, CA, USA) per reaction). For each cDNA sample, a cDNA mix was prepared (1 µL cDNA, 7 µL MQ). qRT-PCR reactions were prepared by combining 12 µL primer mix with 8 µL cDNA mix in a qPCR plate (BIOPlastics, B17489, Landgraaf, the Netherlands). Plates were analyzed in the CFX Thermo Cycler (Bio-Rad), following a three-step program (initial denaturation for 3 min at 95° C, 40 cycles of 95°C for 10 sec, 30 sec at annealing temperature, and 30 seconds at 72°C, followed by a melt curve from 95 to 55° C). Raw data was analyzed with the LinRegPCR software (version 2021.2) and, where necessary, variation between plates was reduced with Factor_qPCR (version 2020.0)^12,13^.

### Statistical analysis

Single and multiple linear regression analyses were run in IBM SPSS Statistics (v28.0.1.1 (15)). Residuals were inspected for homogeneity. Student’s t-tests were conducted in Microsoft Excel (v2410).

## RESULTS

### Correlation between *DAM* expression and semi-dormancy

To induce semi-dormancy, strawberry plants were exposed to short day conditions. To subsequently break dormancy, the plants were subjected to chilling. The cultivars ‘Elsanta’ (chill requirement 1300 h) and ‘Fandango’ (700 h) behaved similarly: In short days, petiole length and leaf area of newly emerged leaves decreased after the start of the experiment and remained low during the induction phase (Figure 4A-D). A similar decrease in petiole length and leaf area was seen in plants in the far-red extended treatment, although the reduction in vegetative growth in this treatment was generally delayed compared to the short day treatment. All ‘Fandango’ plants produced runners during the starting phase as well as some of the ‘Elsanta’ plants. During the induction phase, runner production of both cultivars dropped to zero (Figure 4G-H).

**Figure 4.**
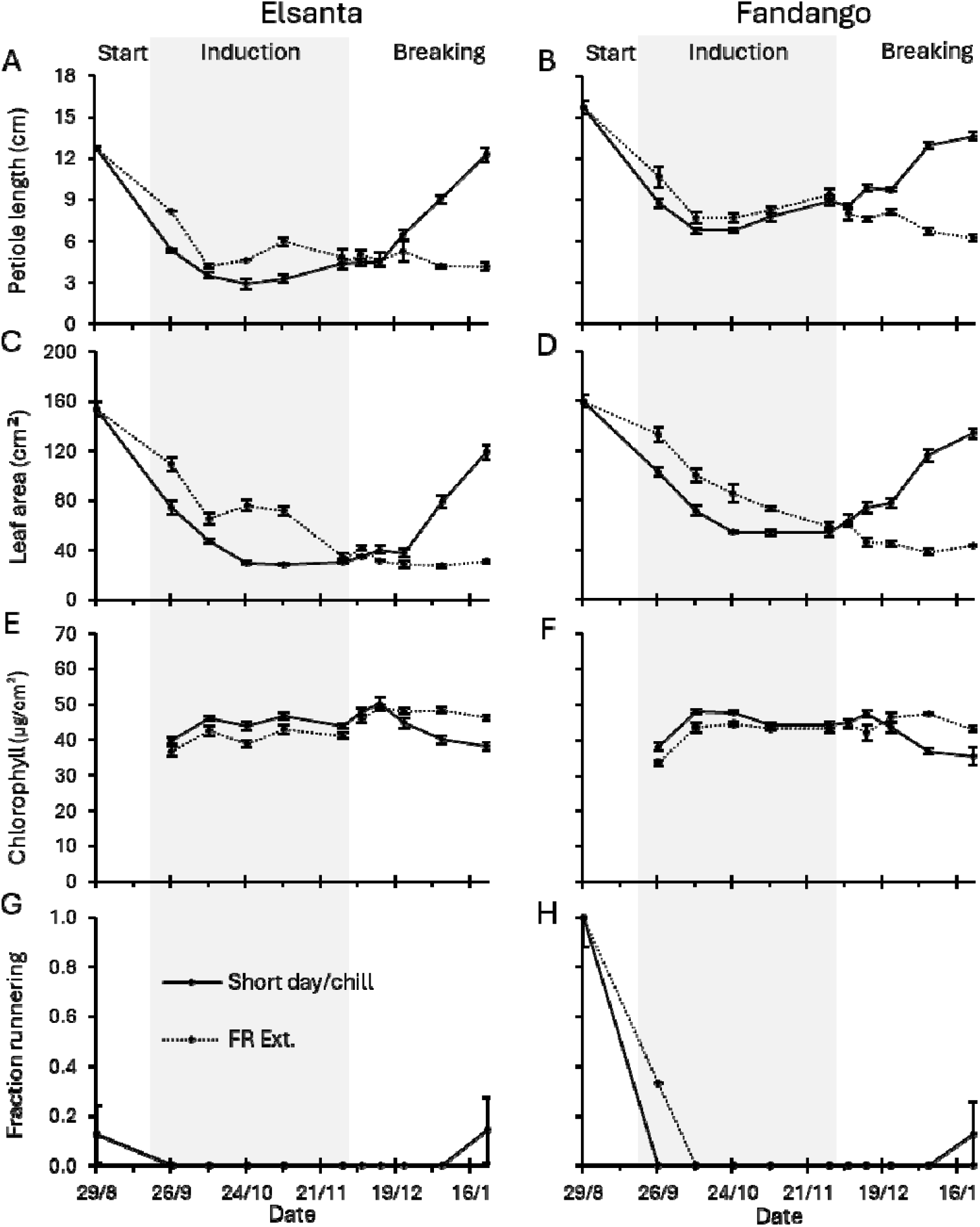
morphological and physiological measurements after four weeks of forcing. During the induction phase (indicated by the grey area), plants were grown in the short day or far-red extended treatment. During the breaking phase plants were chilled (2°C in complete darkness) or grown in the far-red extended treatment. Petiole length, leaf area, and chlorophyll content indicate average of measurements of all newly emerged and fully expanded leaves during forcing. Fraction runnering indicates the fraction of plants that produced at least one runner during forcing. The x-axisdenotes the date on which plants were potted and subsequently forced for four weeks in the far-red extended compartment. N=8, error bars indicate the standard errors.

For both cultivars in chilling conditions, the petiole length and leaf area of newly emerged leaves increased rapidly after the third time point of the breaking phase (Figure 4A-D). In contrast, petiole length and leaf area of plants from far-red extended conditions did not increase. In ‘Fandango’, vegetative growth already increased slightly between the first and third time points of the breaking phase. However, similarly to ‘Elsanta’, the most notable increase in vegetative growth occurred after the third time point of the breaking phase. In both cultivars, leaf chlorophyll content of plants from the chilling treatment decreased after the third time point of the breaking phase, resulting in visually brighter green leaves compared to plants from the far-red extended treatment (Figure 4E-F). On the final time point, some plants of both cultivars in the chilling treatment produced runners, while none of the plants in the far-red extended treatment did so (Figure 4G-H).

‘Elsanta’ that were forced during the starting phase did not flower (Figure 5A). In the induction phase, ‘Elsanta’ plants grown in short day conditions started flowering earlier than the ones grown in far-red extended conditions, which flowered with a few weeks of delay. A similar pattern was seen in ‘Fandango’ (Figure 5B). None of the plants flowered until the third time point of the induction phase, after which the fraction of flowering plants quickly increased. The plants from far-red extended conditions flowered with a slight delay compared to those in short day conditions.

**Figure 5.**
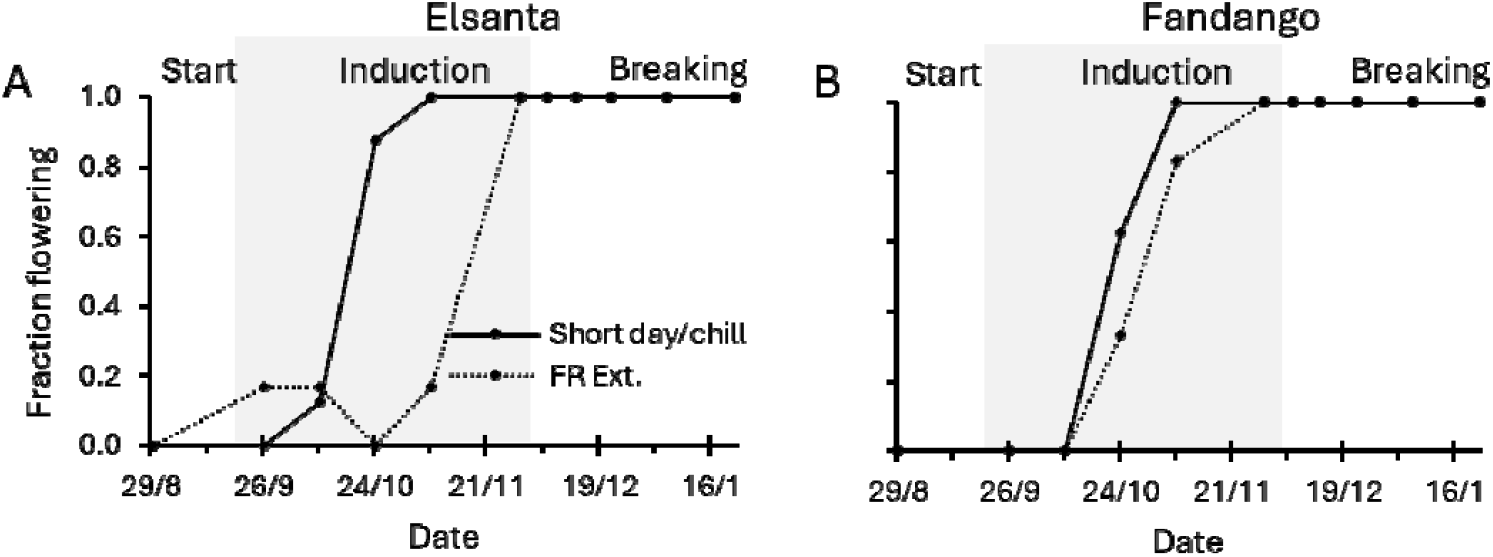
fraction of plants that produced flowers during forcing. During the induction phase (indicated by the grey area), plants were grown in short day or far-red extended conditions. During the breaking phase plants were chilled (2°C in complete darkness) or grown in far-red extended conditions. The x-axis denotes the date on which plants were potted and subsequently forced for four weeks in the far-red extended compartment, after which the flowering plants were counted. Data points denote the fraction of plants that had at least one open flower. Solid lines indicate short day (induction phase) and chilling (breaking phase) treatments, dotted lines indicate far-red extended treatments. N=8.

The number of meristem positions remained stable throughout the induction phase. ‘Elsanta’ plants had slightly more meristem positions than ‘Fandango’ plants (Figure 6A). Throughout most of the induction phase, all plants of both cultivars had a single crown. On the last time point of the induction phase, some plants, of which the majority of the cultivar ‘Fandango’, had produced a second crown (Figure 6B). The percentage of the total meristems that had turned from vegetative to generative increased throughout the induction phase in both cultivars, in short day as well as far-red extended conditions (Figure 6C). In ‘Elsanta’ plants, with a single exception on the second time point, no runner initials were found on any time point in the induction phase. ‘Fandango’ plants did produce runner initials during the induction phase. The number of runner initials that were produced generally decreased over time and reached zero on the last time point (Figure 6D).

**Figure 6.**
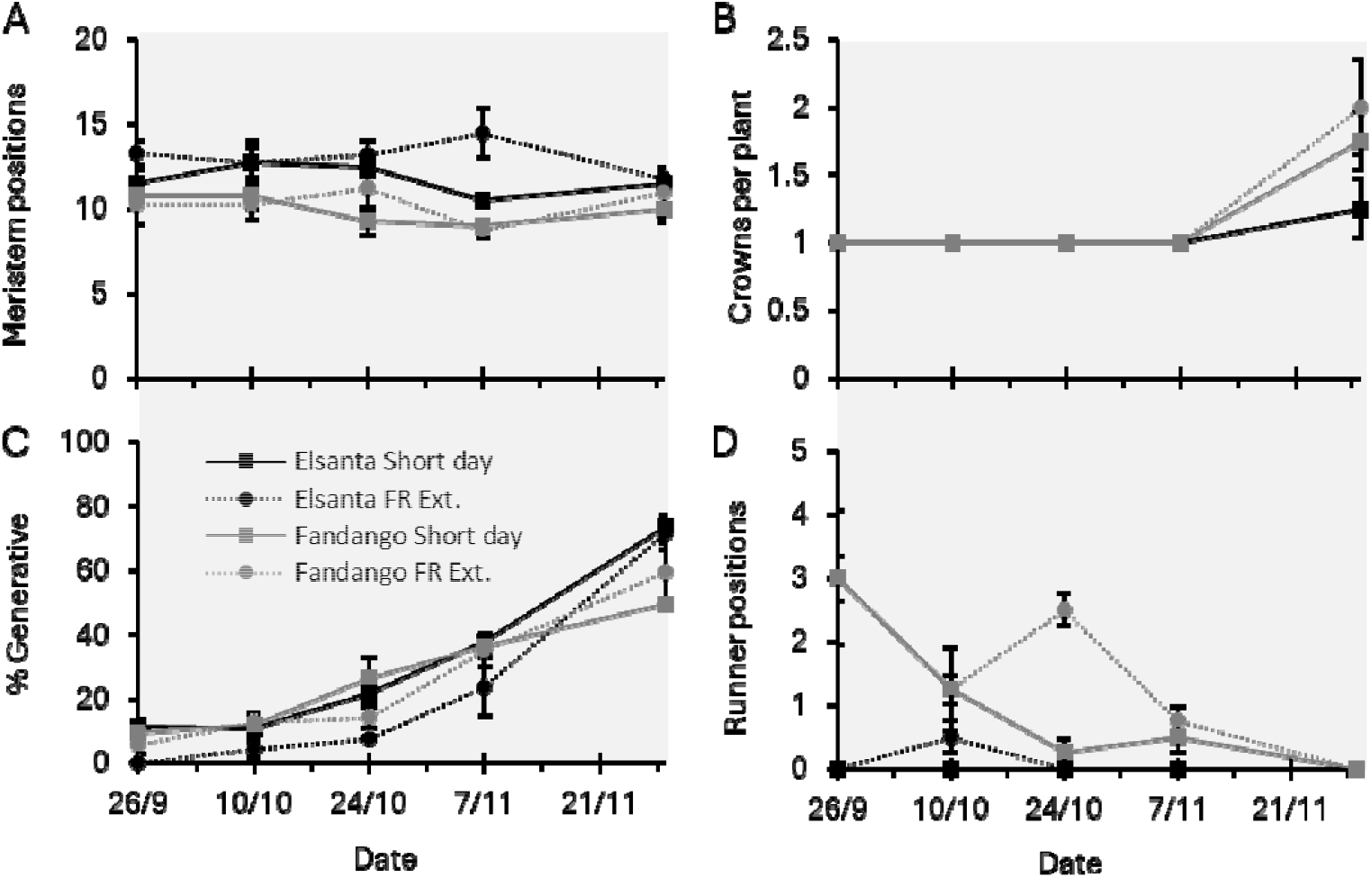
meristem analysis. During the induction phase, plants were grown in short day or far-red extended conditions. On each time point during the induction phase, ‘Elsanta’ and ‘Fandango’ plants were dissected and meristems were analyzed. N=4, error bars indicate the standard errors.

In ‘Elsanta’ as well as ‘Fandango’, the expression of *DAM3* and *DAM4* increased during the induction phase in the short day as well as the far-red extended treatment (Figure 7). During the breaking phase, *DAM3* and *DAM4* expression decreased in both cultivars in the chilling treatment, whereas gene expression of plants in the far-red extended treatment remained stable or further increased.

**Figure 7.**
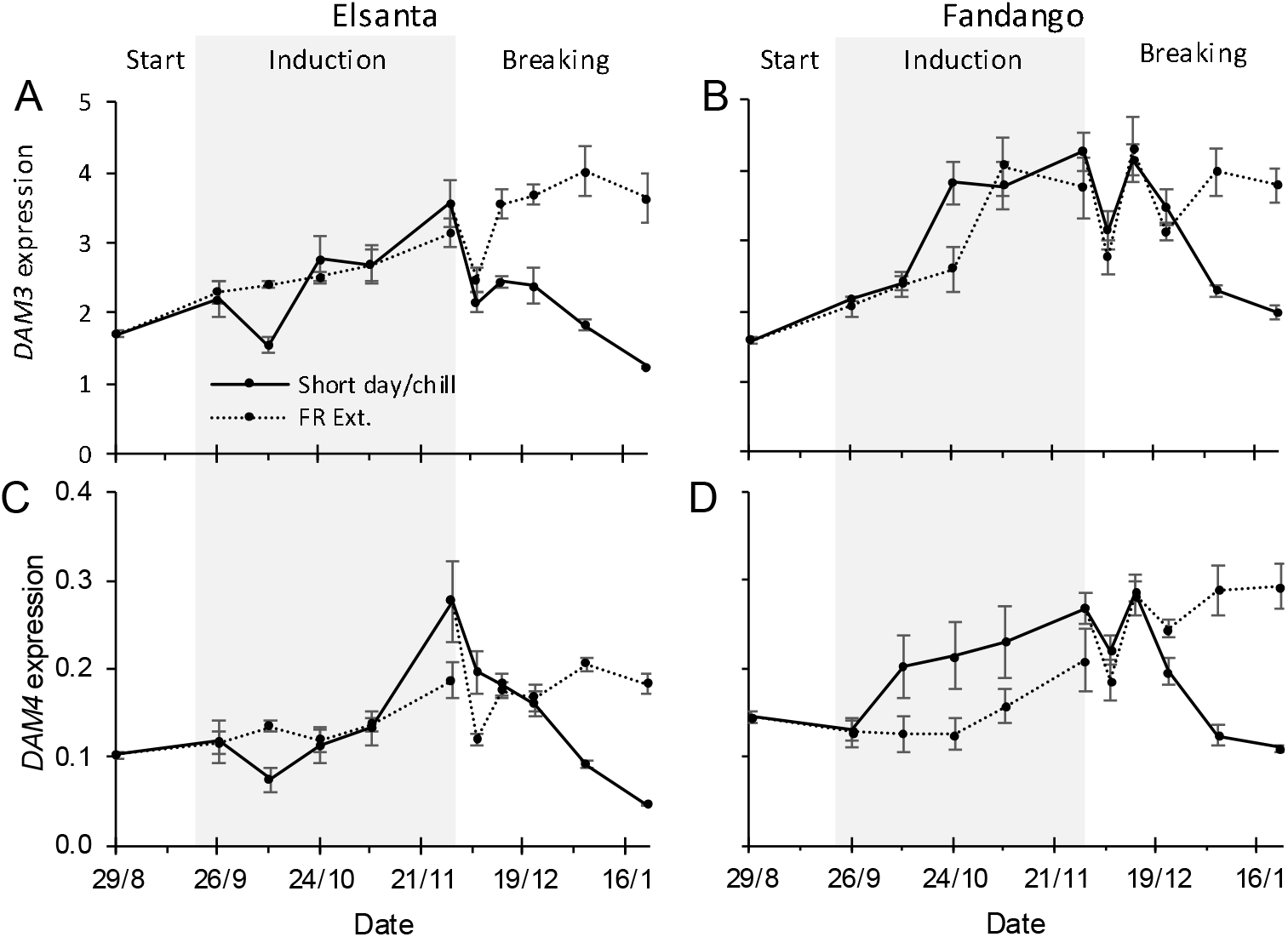
relative expression of *DAM3* and *DAM4* in leaves. During the induction phase (indicated by the grey area), plants were grown in short day or far-red extended conditions. During the breaking phase plants were chilled (2°C in complete darkness) or grown in far-red extended conditions. On each time point, leaf samples were taken from ‘Elsanta’ (left column) and ‘Fandango’ plants (right column). Expression values were determined by qRT-PCR and normalized for the expression of the reference genes *FaDBP, FaMSI*, and *FaACTIN1*. N=4, error bars denote standard errors.

Linear regression was done to analyze the relationship between *DAM* expression and vegetative growth. Ambient temperature was included in the analysis to correct for temperature changes (Table 2). In the dormancy induction phase, the expression of *DAM3* or *DAM4* correlated with vegetative growth in half of the performed analyses. However, the effect of ambient temperature during the forcing period was always significant in the induction phase. *DAM3* was more often found to correlate with vegetative growth than *DAM4*.

During the dormancy breaking phase, ambient temperature did not significantly affect leaf area or petiole length (Table 2). Linear regression between *DAM* expression and vegetative growth was performed on these data points (Table 3). The relative expression of *DAM3* and *DAM4* showed a linear correlation with petiole length and leaf area. DAM4 expression was found to be a better predictor of vegetative growth than DAM3 expression. Scatter plots of the regression data have been included in Supplement 3.

**Table 3.** Regression results of vegetative growth as a function *DAM* expression and temperature. Values were obtained by multiple linear regression of leaf area or petiole length (dependent variables) with *DAM* relative expression and average temperature (T) during the forcing period (independent variables). Asterisks indicate significance: ^*^ P ≤ 0.05, ^**^ P ≤ 0.01, ^***^ P ≤ 0.001. NS = not significant.

| Indep. var | Phase | Dep. var | 'Elsanta' |  |  | 'Fandango' |  |  |
| --- | --- | --- | --- | --- | --- | --- | --- | --- |
|  |  |  | Sig. *DAM* | Sig. Temp | R<sup>2</sup> | Sig. *DAM* | Sig. Temp | R<sup>2</sup> |
| *DAM3* + T | Induction | Leaf Area | * | * | 0.702 | *** | ** | 0.897 |
|  |  | Petiole length | NS | * | 0.843 | * | *** | 0.948 |
|  | Breaking | Leaf Area | * | NS | 0.574 | * | NS | 0.589 |
|  |  | Petiole length | * | NS | 0.583 | * | NS | 0.582 |
| *DAM4* + T | Induction | Leaf Area | NS | ** | 0.674 | *** | ** | 0.850 |
|  |  | Petiole length | NS | *** | 0.832 | NS | *** | 0.936 |
|  | Breaking | Leaf Area | *** | NS | 0.805 | ** | NS | 0.766 |
|  |  | Petiole length | *** | NS | 0.855 | ** | NS | 0.743 |

### *DAM* expression in relation to chill requirement

Expression of *DAM3* and *DAM4* in leaf was upregulated during the induction phase and downregulated during the breaking phase in all four cultivars. For *DAM3*, the peak of expression of the cultivars ‘Favori’ and ‘S2017-0160’ (chill requirement of 1100 and 400 h respectively) was higher than that of ‘Elsanta’ and ‘Fandango’ (1300 and 700 h). The rate of up- and downregulation of *DAM3* relative to the first time point was similar in all four cultivars. The peak of *DAM4* expression was notably lower in ‘Favori’, compared to the other cultivars, but the expression pattern relative to the first time point was highly similar in all cultivars (Figure 8).

**Figure 8.**
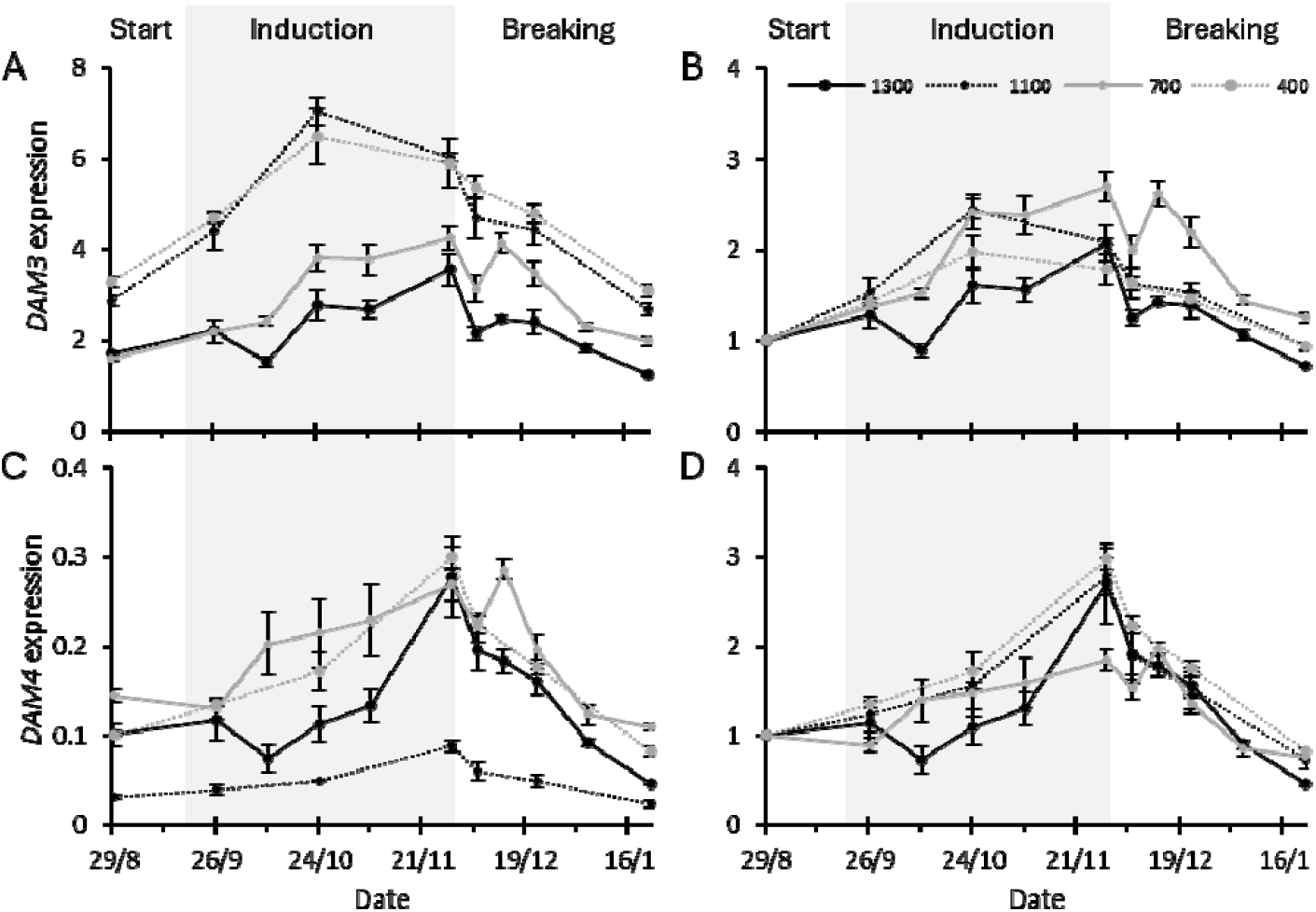
relative expression of *DAM3* and *DAM4* in leaves of cultivars with different chilling requirements. During the induction phase (indicated by the grey area), plants were grown in short day or far-red extended conditions. During the breaking phase plants were chilled (2°C in complete darkness) or grown in far-red extended conditions. Leaf samples were taken on each time-point. Chill requirements are expressed in accumulated hours of chilling below 7° C. 1300: ‘Elsanta’; 1100: ‘Favori’; 700: ‘Fandango’; 400: ‘S2017-0160’. Left column: Expression values were determined by qPCR and normalized for the expression of the reference genes *FaDBP, FaMSI*, and *FaACTIN1*. Right column: the first time point of each cultivar was normalized to 1. N=4, error bars denote standard errors.

### *DAM* expression between different leaf types

At the end of the forcing period, *DAM* expression was compared between winter leaves (emerged before chilling) and summer leaves (emerged after chilling, during the forcing period). The relative expression of *DAM4* in ‘Elsanta’ was lower in summer leaves than in winter leaves (Figure 9G). However, there was no difference in gene expression pattern over time between the two leaf types. No clear differences in gene expression level or pattern over time were found between the two leaf types for the other *DAM* genes in either cultivar (Figure 9A-F,H).

**Figure 9.**
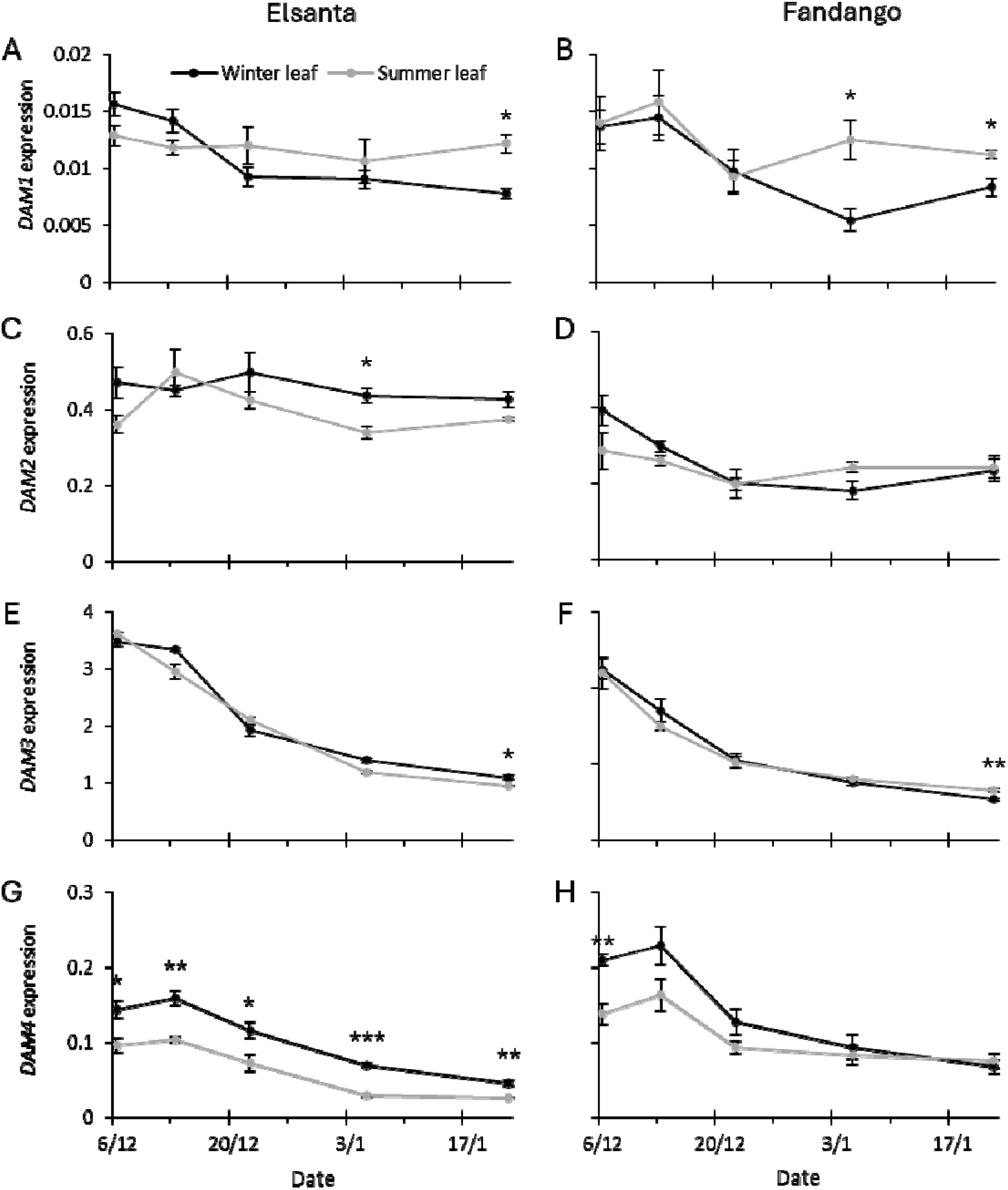
Relative expression of *DAM1, DAM2, DAM3*, and *DAM4* in winter and summer leaves of ‘Elsanta’ and ‘Fandango’. Winter leaves are leaves that emerged before the plants were exposed to chilling, summer leaves emerged during the forcing period after the chilling treatment. Dates on the x-axis denote the start of the four week forcing period. T-test was used to analyze differences in expression between leaf types. Asterisks indicate significance: * P ≤ 0.05, ** P ≤ 0.01, *** P ≤ 0.001.

## DISCUSSION

Earlier work pointed at *DAM3* and *DAM4* as potential regulators of dormancy in strawberry^10^. In the current work, *DAM3* and *DAM4* expression, plant growth and development during dormancy induction and breaking were analyzed in detail to further elucidate the involvement of these genes in dormancy regulation. The correlation between *DAM* expression and chill requirement was analyzed by comparison of *DAM3* and *DAM4* expression between cultivars with different chill requirements. We tested whether there are differences in *DAM* expression between leaf types by comparing gene expression between winter and summer leaves.

### Short day and far-red extended conditions induced semi-dormancy

The analysis of plant morphology and development suggests that, by the end of the dormancy induction phase, plants in short day as well as far-red extended conditions were in a similar state of semi-dormancy. The far-red extended treatment was included to show that the changes in plant morphology, physiology and gene expression in the induction phase were caused by exposure to short day conditions, thereby excluding other factors, such as plant age or ambient temperature.

We observed a decrease in petiole length and leaf area of newly grown leaves during the dormancy induction phase in plants exposed to short day conditions (Figure 4A-D). This decrease in vegetative growth is consistent with the induction of semi-dormancy and the transition to the winter morphology^3^. Although slightly delayed, the decrease in vegetative growth of plants in far-red extended conditions was comparable to that of the plants in short day conditions. This is corroborated by the decrease in runner formation (Figure 4G, H), the increase in flower emergence (Figure 5), and the increase in crowns and generative meristem during the induction phase (Figure 6B, C), which were observed in both growing conditions. Together, these results point to a short day developmental response which is linked to the induction of semi-dormancy^2,9^.

Besides short day conditions, generative development in strawberry can be induced by exposure to sufficiently low temperatures, which overrides the photoperiodic effect^17,18^. However, in *F. ananassa* cultivar Korona, a 16 hour photoperiod was reported not to result in generative growth at a day temperature of 12°C, and ‘Elsanta’ plants in long day conditions did not flower after being exposed to 9°C for over a month^19,20^. In our experiment, average daily temperatures (combining day and night) of 13° C or lower did not occur until October 25 (Figure 3A). Meristem analysis showed that flower initiation in both conditions had already occurred by October 24 (Figure 6). It is therefore more likely that the daylength extension with far-red was not sufficient to elicit a long day response of the plants, rather than that low temperatures induced generative growth. Lamps emitting a mixture of white, red, and far-red light were used for extension of the photoperiod. A potential reason that a long day response was not observed in the plants in these conditions is that the light intensity of the day length extension was too low to elicit this response.

### Chilling broke semi-dormancy during the breaking phase

During the dormancy breaking phase, extended exposure to cold temperatures, restored petiole length and leaf area to levels similar to the start of the experiment, indicating semi-dormancy breaking (Figure 4A-D)^2,4^. Although, similar to the induction, the breaking of semi-dormancy shows a quantitative response, this response only initiated after an initial accumulation of 336 to 552 chill hours. Below this amount, there were no or only minimal effects of chilling on plant morphology. We also observed gradually decreasing chlorophyll content in the leaves of plants that received the chilling treatment, compared to those in the far-red extended treatment, consistent with the transition of winter morphology to summer morphology that accompanies the breaking of semi-dormancy^3^.

### *DAM3* and *DAM4* are downregulated by chilling

Chilling reduced *DAM3* and *DAM4* expression during the breaking phase (Figure 7). The expression levels did not change or even increased in plants in far-red extended conditions. The drop in *DAM3* and *DAM4* expression after accumulation of more than 552 chill hours, and the concurrent increase in vegetative growth, coincides with accumulated amount of chill hours after which flower initiation under short day conditions is no longer possible (T. van Dijk, Fresh Forward, personal communication). This suggests that high expression of *DAM3* and/or *DAM4* may be a requirement for flower initiation. A similar effect was described by Sønsteby and Heide (2006), who reported that avoidance of the winter morphology, by exposing the plants to temperatures of 6°C or lower while in short day conditions, simultaneously inhibited flower initiation^2^. These results suggest that *DAM3* and *DAM4* do not only regulate winter morphology, but rather act as a switch between vegetative and generative development. In peach, a natural knock-out of several *DAM* genes resulted in an *evergrowing* phenotype which maintained bud growth in conditions that are otherwise inductive for semi-dormancy^7,8^. In strawberry, the loss of *DAM3* and *DAM4* expression might result in the inability to initiate flowers under short day conditions.

### *DAM4* expression explains vegetative growth during semi-dormancy breaking

*DAM4* expression explained a high amount of variation in vegetative growth (up to ~85%) which is consistent with its hypothesized role as master regulator of semi-dormancy (Table 3, Table 4). The higher variability in ambient temperature during the induction phase (Figure 3A), may explain the lack of correlation between *DAM* expression and vegetative growth in this phase of the experiment. This suggests that the effect of temperature on vegetative growth overrides that of *DAM* expression. Moreover, is it possible that the decreased correlation between *DAM* expression and vegetative growth during the induction phase has a plant developmental reason, but this cannot be inferred from the current dataset. In the dormancy breaking phase, the high significance and R-squared values of *DAM4* compared to *DAM3* indicate that DAM4 expression is a better predictor of vegetative growth than DAM3 expression. This is somewhat surprising given that the expression level of DAM4 is roughly an order or magnitude lower than that of DAM3.

**Table 4.** Regression results of vegetative growth as a function *DAM* expression. Values were obtained by linear regression of leaf area or petiole length (dependent variable) versus *DAM* relative expression (independent variable). Cells are colored on a scale from blue to red according the R-squared value from low to high. Asterisks indicate significance of single linear regressions: ^*^ P ≤ 0.05, ^**^ P ≤ 0.01, ^***^ P ≤ 0.001z

| Indep. var | Phase | Dep. var | 'Elsanta' |  | 'Fandango' |  |
| --- | --- | --- | --- | --- | --- | --- |
|  |  |  | Sig. *DAM* | R<sup>2</sup> | Sig. *DAM* | R<sup>2</sup> |
| *DAM3* | Breaking | Leaf area | * | 0.569 | ** | 0.585 |
|  |  | Petiole length | * | 0.564 | * | 0.533 |
| *DAM4* |  | Leaf area | *** | 0.793 | *** | 0.766 |
|  |  | Petiole length | *** | 0.845 | *** | 0.720 |

### Expression of *DAM3* and *DAM4* does not correlate with chill requirement

Analysis of *DAM* gene expression in multiple peach (*Prunus persica*) cultivars, suggested a correlation between the expression of *PpDAM5* and *PpDAM6* and the chilling requirement of a cultivar during chilling accumulation^21^. We found differences between cultivars in peak expression of *DAM3* and *DAM4* during the dormancy induction phase (Figure 8). However, these differences did not correlate with the chill requirement of the cultivars. In addition, the differences in expression were not consistent between the two genes. We also did not observe differences in the rate of gene up- or downregulation during induction and breaking respectively. Taken together, the data suggest that the chill requirement is not a function of *DAM3* or *DAM4* expression.

The cultivar ‘Fandango’ is reported to need fewer chilling hours than the cultivar ‘Elsanta’, which is why they are referred to as low-chill and high-chill cultivars respectively. However, morphological measurements as well as gene expression analysis suggest that both cultivars require in excess of 552 hours at 2° C to substantially reduce *DAM* expression and increase vegetative growth. We found that for a certain amount of exposure to short days or chilling, ‘Fandango’ plants always had larger leaves and longer petioles than ‘Elsanta’ plants. For example: after accumulation of 336 chilling hours, the petiole length of ‘Fandango’ was 9.9 cm; after chilling for 888 hours, the petiole length of ‘Elsanta’ was only 9.1 cm (Figure 4A-B). Without the context of how vegetative growth behaves at different amounts of chilling, ‘Fandango’ plants appear to need less cold exposure to break semi-dormancy, while in reality ‘Fandango’ has increased vegetative growth (at least pertaining to petiole length and leaf area) compared to ‘Elsanta’, regardless of dormancy status. Our findings suggest that both cultivars require over 552 hours of chilling exposure to break semi-dormancy, and around 1296 hours to fully restore vegetative growth.

### No difference in *DAM* expression was found between winter and summer leaves

In an earlier work, differential expression of *DAM1* and *DAM2* was found between summer and winter leaves after five weeks of forcing (at 1140 accumulated chill hours), suggesting a role for these genes in the developmental regulation of these leaf types^10^. In the current study we expanded on this analysis by including multiple time points with increasing amounts of accumulated chilling hours. Because increasing the amount of chilling hours leads to increased morphological differences between the leaf types (Figure 4, breaking phase), we would expect the difference in gene expression between these leaf types to increase concurrently. However, in the current study no such differences were found. The relative expression of *DAM1* and *DAM2* fluctuated, which resulted in differential expression between winter and summer leaves at some time points (Figure 9), which may explain the preliminary results.

*DAM4* expression in ‘Elsanta’ was found to be consistently lower in summer leaves compared to winter leaves. This underlines the importance of consistency in the selection of leaf types during expression analysis of *DAM* genes, since alternating between leaf types might lead to perceived changes in expression level and pattern. The pattern of expression over time was found to be highly similar between leaf types for *DAM3* and *DAM4*, which suggests that any leaf type can be used to comment on *DAM* expression in the leaves of the plant as a whole.

## CONCLUSIONS

We conclude that the expression of *DAM3* and *DAM4* negatively correlates with vegetative growth during the breaking of semi-dormancy. The expression pattern of these genes is consistent with their hypothesized role as regulators of semi-dormancy and the switch between vegetative and generative development. The chill requirement of strawberry cultivars does not correlate with *DAM3* or *DAM4* expression level or pattern, and likely is a measure of vegetative growth independent from dormancy status.

## ACKNOWLEDGEMENTS

This research is part of the TTW Perspectief programme “Sky High”, which is supported by AMS Institute, Bayer, Bosman van Zaal, Certhon, Fresh Forward, Grodan, Growy, Own Greens/Vitroplus, Priva, Signify, Solynta, Unilever, van Bergen Kolpa Architects, and the Dutch Research Council (NWO).

## DATA AVAILABILITY STATEMENT

The morphological, physiological and flowering data (Figure 4 and 5), meristem data (Figures 6), and gene expression data (Figures 7, 8 and 9) are publicly available in the figshare repository: https://doi.org/10.6084/m9.figshare.28981544.

