## Supplement for "*Fragaria ananassa DAM4* expression correlates with vegetative growth during semi-dormancy breaking"

### Supplement 1

**SON-T, spectrum.**

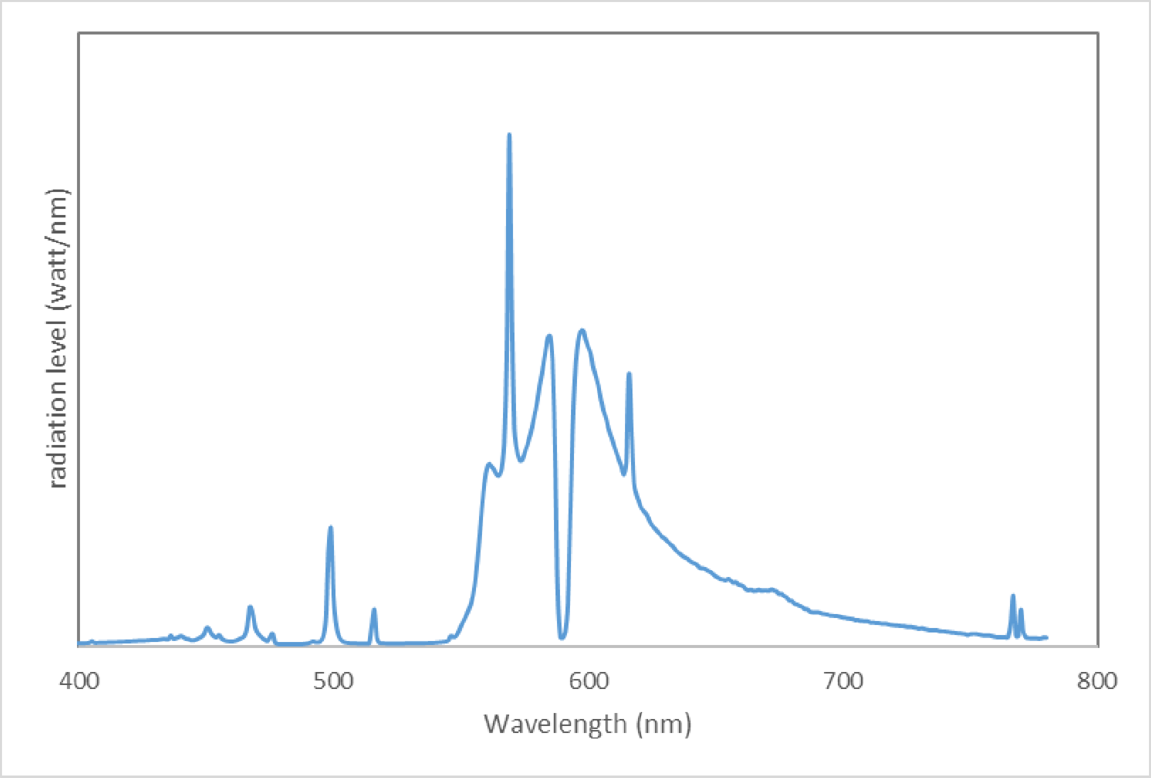

### Supplement 2

**GreenPower LED Flowering Lamp Gen. 2.1 (DRWFR), spectrum.**

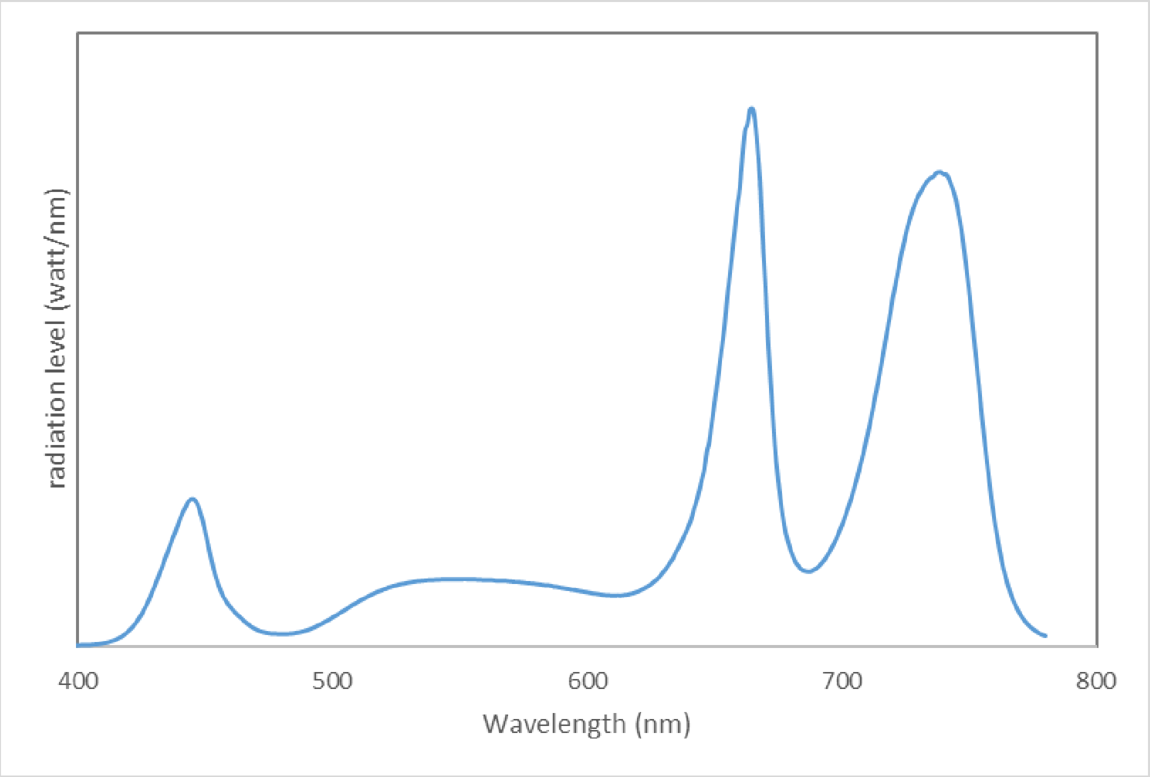

### Supplement 3

**Regression of *DAM3* and *DAM4* relative expression with leaf area and petiole length.** Regression of *DAM3* and *DAM4* relative expression with leaf area and petiole length. The data of the start and induction phase (black) and the breaking phase (blue) were analyzed separately. Asterisks following the $R^2$ values indicate significance: * P ≤ 0.05, ** P ≤ 0.01, *** P ≤ 0.001

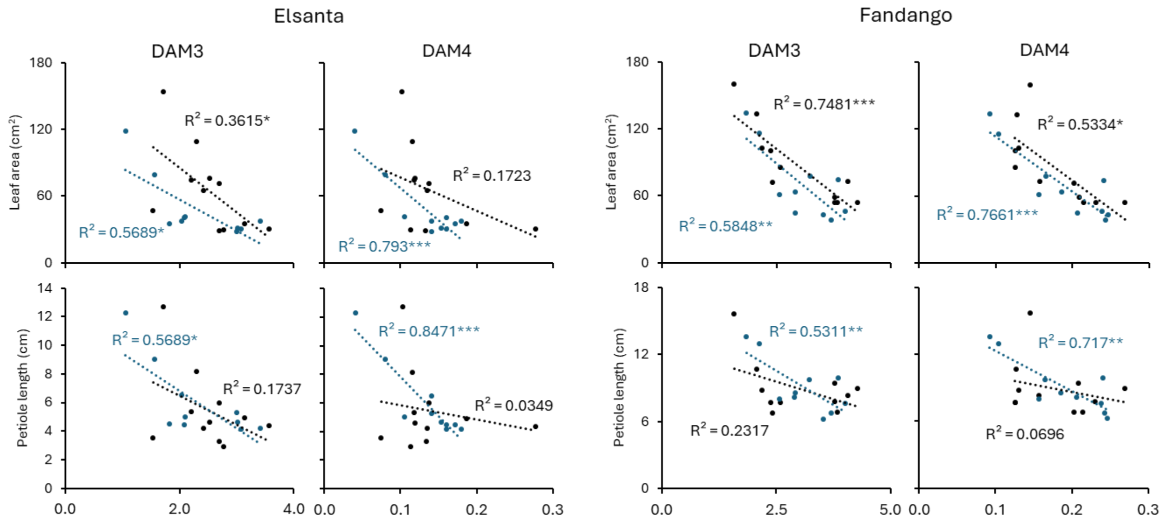
